# Significant Shortest Paths For The Detection Of Putative Disease Modules

**DOI:** 10.1101/2020.04.01.019844

**Authors:** Daniele Pepe

**Author notes:** Email address: DP.

## Abstract

**Background:** The characterization of diseases in terms of perturbated gene modules was recently introduced for the analysis of gene expression data. Some approaches were proposed in literature, but many times they are inductive approaches. This means that starting directly from data, they try to infer key gene networks potentially associated to the biological phenomenon studied. However they ignore the biological information already available to characterize the gene modules. Here we propose the detection of perturbed gene modules using the combination of data driven and hypothesis-driven approaches relying on biological metabolic pathways and significant shortest paths tested by structural equation modeling.

The procedure was tested on microarray experiments relative to infliximab response in patients with inflammatory bowel disease. Starting from differentially expressed genes (DEGs) and pathway analysis, significant shortest paths between DEGs were found and merged together. The validation of the final disease module was principally done by the comparison of genes in the module with those already associated with the disease, using the Wang similarity semantic index, and enrichment analysis based on Disease Ontology. Finally a topological analysis of the module via centrality measures and the identification of the cut vertices, allowed to unveil important nodes in the network as the TNF gene, and other potential drug target genes as p65 and PTPN6.

**Conclusions:** Here we propose a downstream method for the characterization of disease modules from gene expression data. The core of the method is rooted on the identification of significant shortest paths between DEGs by structural equation modeling. This allows to have a mix approach based on data and biological knowledge enclosed in biological pathways. Other methods here described as enrichment analysis and topological analysis were functional to the validation of the procedure. The results obtained were promising, considering the genes and their connections found in the putative disease modules.

## Background

The reductionist approach in medicine, based on the principle “of divide and conquer”, although useful, presents limits when it is necessary to explain the onset and progression of complex diseases. In fact the approach is rooted in the assumption that if a complex problem is divided into more understandable and smaller units, then by their reconstruction, it is possible to unveil the complex problem treated. For this reason, there are lists of genes associated with diseases. OMIM, a free database [1], for example, offers a catalogue of genes with the relative description of their role in the associated phenotypes. Conversely to this point of view, in the 1972, Anderson, in the article “More is complex” [2], affirms that the behavior of large and complex aggregate of elementary particles cannot be understood in terms of a simple extrapolation of the properties of a few particle. At each level of complexity entirely new properties appear. For this reason, we are assisting to the passage from the reductionist approach to the systemic approach [3]. Most of biological networks are subjected to specific laws [4], as the small world phenomena, which affirms that there are relatively short paths between any pair of nodes, the scale-free principle, with the consequence that there are few highly connected hubs; the local hypothesis i.e. the presence of modules, highly interlinked local regions in the network in which the components are involved in same biological processes. The analysis of high-throughput biological data as the microarray gene expression data for many years was influenced by the reductionist approach. The goal of the analysis was to supply the signature of a disease, i.e., the list of differentially expressed genes (DEGs). These methods tend to ignore much of the signal in the data, both in genes whose activity changes but does not pass the threshold for differential expression and in genes that are differentially expressed but unfamiliar to the researcher analyzing the list. Listing genes associated with a certain disease is far from identifying the biological processes in which these genes are involved. Developing methods that can supply more biological information in terms of connections between DEGs could allow to understand the key processes giving rise to the disease. There is an increasing evidence that a functional gene system is composed of coordinated and independent modules. Modularity is a general design principle in biological systems and has been observed also in transcriptional regulation networks [5]. Applying module level analysis should help to study biological systems at different levels and to understand which properties characterize the level of complexity considered. Many approaches exist that use a gene-module view as the basic building blocks of the analysis [6]. In general, it is used to divide the module identification in three main approaches: 1) network-based approach; 2) expression-based approach and 3) pathway-based approach [7].

The first approach is based on the topology of network, and modules are defined as subsets of vertices with high connectivity between them and less with external nodes. The second approach uses gene expression data for inferring modules of genes exhibiting similar expression by, for example, clustering methods. The third approach detects expression changes in biological pathways, group of genes that accomplishes specific biological functions.

The approach proposed in this paper, it is a mix and more general approach that takes advantage of the three approaches previously described. In fact the pathway-based approach was used to detect perturbed KEGG pathways [8], then the network-approach was employed to identify the shortest paths between the DEGs and finally significant shortest path (SSPs) s were found using the expression data and structural equation modeling (SEM) [9]. The idea to consider shortest paths between DEGs to understand how they are connected was already applied in the pipeline proposed by Pepe & Grassi [10]. However, differently from the method proposed by them, each shortest path is obtained by the network obtained by the fusion of the significant pathways. Furthermore each shortest path is considered a structural model and tested by multiple group SEM. All the SSPs were joined to have the final perturbed disease module.

## Methods

The classical differential gene expression analysis allows to identify DEGs. In our analysis we used the t-test with Benjamini-Hochberg (BH) false discovery rate correction, but any other procedure can be used. The DEGs was used to find the corresponding perturbed pathways using the “Signaling Impact Pathway Analysis” (SPIA) [11]. This algorithm takes in consideration two evidences for the detection of perturbed pathways: enrichment analysis and perturbation analysis. The corresponding perturbed KEGG pathways can be represented as mixed graphs, where the nodes represent genes and the edges represent multiple functional relationships between genes as activation, inhibition, binding etc. The core idea for the building of a disease module is to understand how the DEGs in the perturbed pathways are connected between them. The first step for reaching this goal is to merge all the significant pathways in a unique graph. The second step is to find the significant shortest paths that put in communication every couple of DEGs. Each shortest path could be represented as a list of nodes P*=*(*p*_*i*_, *p*_*i+1*_,..,., *p*_*j-1*,_ *p*_*j*_) and a list of the corresponding edges E=(e_i(i+1)_, …, e_(j-1)j_) where (*p*_*i*_, *p*_*j*_) are DEGs and*∈ DEG* (*p*_*i+1*_,…, *p*_*j-1*_) can be DEGs or other microarray genes. In our case every edge is directed. A shortest path can be codified as a structural equation (SE) model, in the following way:

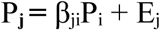

where P_j_ represents every gene in the path that is influenced directly by the gene P_i;_ β_ji_ is the strength of relationship between node P_i_ and P_j_; E_j_ is a term that represent external causes that have an effect on P_j_ but not explicated in the model. Considering that the shortest paths selected are induced paths, every shortest path can be represented by j-1 simple linear equations, where j is the number of nodes in the path.

For the estimation of the parameters β_ij_, the Maximum Likelihood estimation (MLE) is used, assuming that all observed variables have a multinormal distribution. For finding the SSPs, the following omnibus test was performed:

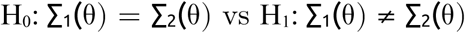

where ∑_1_(θ) and ∑_2_(θ) are the model-implied covariance matrices of the two biological groups, and θ represents the parameters of the model. The statistical significance is determined by comparison of likelihood ratio test (LRT) chi-square values. If there is a significant difference (P<0.05), after the Benjamin-Hochberg correction, in the chi-squared goodness-of-fit index, the shortest path is considered statistically significant. To note that this procedure is a simplified application of SEM as ignoring the quality of fitting, and the respecification of the models. The reason for this choice is that, what we want to know is if the shortest paths, extracted by biological pathways, are significant or not between the groups analyzed. All the SSPs were merged to obtain the final disease module. If the experimental groups are constituted by subgroups, as in the examples showed in this article, an hierarchical analysis is possible. In other words, starting from the set of significant shortest paths, same analysis can be performed but among the subgroups that characterize the experimental conditions. To validate the procedure two different approaches were employed: 1) enrichment analysis based on Disease Ontology (DO) to verify if the module genes are associated to the family of diseases connected with the disease analyzed; 2) semantic similarity index, based on Gene Ontology (GO) terms, between the list of genes associated *“a priori”* to the disease and the list of genes present in the module. DO creates a single structure for the classification of disease and permits to represent them in a relational ontology [12]. For the semantic similarity, the graph-based strategy proposed by Wang et al. [13] was used. The method encodes a GO term’s semantics into a numeric value assigning a value to the contributions of the ancestor terms in the GO graph. Based on the similarity between the GO terms it is possible to measure the functional similarity between list of genes [14]. The similarity goes from 0, when the lists of genes are not connected, to 1, when the lists of genes are associated to the same GO terms. For finding the *a priori* genes a research on Entrez Gene [15] was done, putting the name of the disease and selecting the results only for “Homo sapiens”. This measure encodes the biological meanings associated to a GO term as an aggregation of the semantic contributions of the ancestor terms of the GO term considered. For each set of genes an enrichment analysis on GO biological process (BP), molecular function (MF) and cellular component (CC) terms was done and the list of enriched terms are compared with those obtained by gene disease module by the Wang similarity measure. Finally a basic network analysis was performed based on measures of centralities, as the betweenness and the Bonacich power centrality score, and on the connectivity of the module by the detection of the articulation nodes. The goal of this analysis is to propose genes that can be important for the perturbation of the module on the basis of the network topology. The betweenness centrality for the gene *v* is defined as:

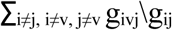

where *g*_*ivj*_ are the shortest paths between the node *i* and *j* in which the node *v* is present, while *g*_*ij*_ is the number of all shortest paths between the nodes *i* and *j*. The normalized betweenness centrality is obtained:

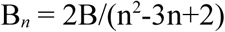

where B is the raw betweenness and n the number of nodes.

The Bonacich power measure corresponds to the notion that the power of a vertex is recursively defined by the sum of the power of its alters. The formula is the following:

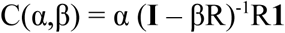

where α is a scaling vector, β is an attenuation factor to weight the centrality of the nodes, R is the adjacency matrix, **I** is the identity matrix, **1** is a matrix of all ones. The articulation nodes are the minimum set of nodes which removal increases the number of connected components. So they represent the nodes that allow to have a connected module.

The procedure was tested on the dataset GSE16879 finalized to identify mucosal gene signatures predictive of response to infliximab in patients with inflammatory bowel disease (IBD) and to gain more insight into the pathogenesis of IBD. The experiment is composed by 133 samples in which different groups are present. The samples represent whole genome gene expression microarray profiles (Affymetrix Human Genome U133 Plus 2.0 Arrays) from total RNA isolated from endoscopic-derived intestinal mucosal biopsies obtained from IBD patients and healthy (non-IBD) controls. For this study, we chose the comparison between 19 samples relative to mucosal biopsies of 19 patients affected by Crohn’s colitis disease (CD) (12 responders and 7 non-responders to infliximab treatment) against 6 controls and the comparison between 24 samples affected by ulcerative colitis (UC) (16 non-responders and 8 responders to infliximab treatment) against 6 controls to verify the goodness of our procedure. It is necessary to highlight that the goal of this paper is to find a way to detect disease modules from perturbed pathways, trying to overcome the limit of the most pathway analysis methods that consider the pathways as independent unities and to detect the key region, in the perturbed network, that could explain the mechanisms of the development of the disease. So it is a downstream analysis performed after the differential analysis and the pathway analysis. The researcher can choose different upstream analysis approaches. Here we used a classical t-test with BH correction and SPIA for pathway analysis, but other choices are possible. All the analyses were performed in R 2.15.3 [16].

## Results and discussion

### CD data

For the CD versus control samples, we found 1245 significant genes using as criteria a BH corrected p-value less than 0.01 and a minimum fold change of 1.5. Starting from DEGs the corresponding module was generated.

#### 3.1 Module generation

On the 1245 DEGs, SPIA revealed important perturbed pathways as the cytokine-cytokine receptor interaction, the chemokine signaling pathway [17] and the NF-▮B signaling pathway. Chronic intestinal inflammation can result from defective immunosuppression, hyper active Th1 cell function or cytokine hyper secretion [18]. Chemokines are excellent candidates for being primarily responsible for the upregulation, perpetuation, and exacerbation of inflammatory and tissue-destructive processes involved in the immunopathogenesis of IBD.[19]. NF-κB pathway regulatesB pathway regulates host inflammatory, immune responses and cellular growth properties by increasing the expression of specific cellular genes. These include genes belonging to the chemokine and cytokine pathways (as *ZAP70* or *PKC*). This pathway, giving the key role in the pathogenesis of inflammatory diseases, was proposed as possible drug target [20]. Giving a look to the direction of the perturbation, the most of the perturbed pathways resulted activated (see Table 1).

**Table 1.** The KEGG perturbed pathways involved in CD. pSize represents the total number of nodes in the pathways, NDE the number of DEGs, tA the total perturbation, pPERT the p-value of the perturbation and pGFdr the adjusted p-value.

| **Name** | **pSize** | **NDE** | **tA** | **pPERT** | **pGFdr** |
| --- | --- | --- | --- | --- | --- |
| Cytokine-cytokine receptor | 241 | 27 | 29.78 | 0.00 | 0.00 |
| Chemokine signaling path. | 178 | 26 | 61.75 | 0.00 | 0.00 |
| Cell adhesion molecules | 88 | 18 | 44.17 | 0.05 | 0.00 |
| B cell receptor signaling path. | 73 | 16 | 10.80 | 0.35 | 0.00 |
| Apoptosis | 84 | 16 | 5.15 | 0.68 | 0.01 |
| Natural killer cell | 128 | 18 | 59.10 | 0.04 | 0.01 |
| HTLV-I infection | 198 | 26 | 17.26 | 0.24 | 0.01 |
| Tuberculosis | 161 | 23 | 13.76 | 0.37 | 0.01 |
| NF-kappa B signaling path. | 75 | 13 | 18.89 | 0.09 | 0.01 |
| Legionellosis | 39 | 9 | 11.50 | 0.11 | 0.01 |
| NOD-like receptor path. | 48 | 10 | 14.40 | 0.10 | 0.01 |
| T cell receptor signaling path. | 97 | 16 | 5.81 | 0.57 | 0.01 |
| Fc epsilon RI signaling path. | 58 | 12 | -1.16 | 0.94 | 0.01 |
| Amoebiasis | 46 | 8 | 12.43 | 0.04 | 0.02 |
| MAPK signaling path. | 252 | 26 | 15.41 | 0.08 | 0.03 |
| Toxoplasmosis | 91 | 13 | 12.03 | 0.20 | 0.04 |
| Jak-STAT signaling path. | 99 | 15 | 0.17 | 0.95 | 0.04 |
| Malaria | 11 | 4 | 2.70 | 0.26 | 0.04 |
| Focal adhesion | 203 | 22 | 20.99 | 0.20 | 0.05 |

This result appears reasonable, considering that a lot of genes, in the perturbed pathways, increase the expression for generating an inflammatory process. For example it is known that *in vivo* expression of all individual chemokines was significantly up-regulated in CD patients when compared with normal controls [21]. The pathways that presents the highest value of active perturbation was the cell adhesion molecules (CAMs). These molecules mediate the leukocyte trafficking from the blood circulation into the sites of inflammation. Not surprisingly, the effects of many anti-inflammatory drugs are the inhibition of the expression of CAMs [22-24]. The pathways were transformed in graph and subsequently merged. This step is important because it permits to consider and connect pathways in a molecular network. This approach allowed to have a more global picture of the molecular basis of the phenomena analyzed. The next step was to find the shortest paths between every couple of DEGs on the total graph. The total number of shortest paths resulted of 7711 with 267 genes involved. For each shortest path a structural model was generated and tested to detect those significant. 567 out of 7711 were found relevant involving 200 genes. Figure 1 shows the graph obtained by the fusion of the 567 significant paths.

**Fig. 1.**
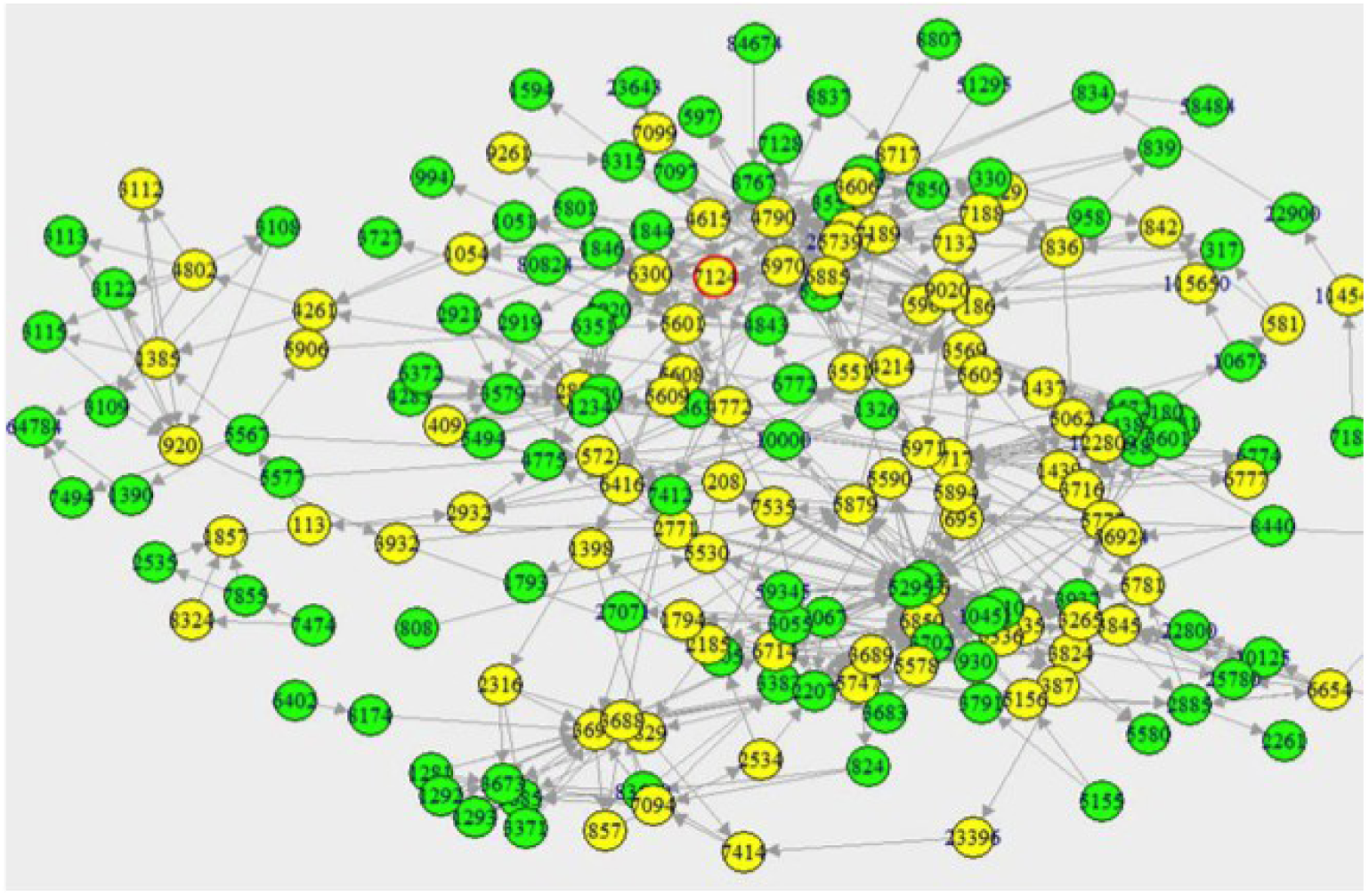
Module obtained by merging the significant shortest paths between DEGs found for CD samples. Green nodes are DEGs while yellow are not DEGs. To note is the gene 7124 circled in red. Notwithstanding this gene was not found as DEG, it plays a fundamental role in the CD.

#### 3.2 Module validation

For the validation of the module two different analyses were performed: 1) one looks for diseases enrichment analysis; 2) the other measures the semantic similarity between genes associated “a priori” with CD with those in the module.

The results of the enrichment analysis on DO was very interesting considering that there are a lot of diseases that share a common root with CD as UC, Rheumatoid arthritis (RA), and Systemic lupus erythematosus (SLE). (see Figure 2). In fact it is known that these diseases share genetic and phenotypic effects with CD [25-28]. Also other diseases as Alzheimer’s disease and autoimmune diseases could be associated with CD, considering that there could be an autoimmune component in the pathogenesis of CD [29]. Finally, it is worthy also to mention the possible co-existence between CD and intestinal endometriosis [30]. For the validation with the semantic similarity approach, we looked for the genes associated with CD in the database “Gene” of NCBI. We found 446 unique Entrez identifiers. The intersection of the genes in the module with those associated to CD revealed 28 genes in common. Of these only 9 were DEGs while the remaining genes were not DEGs. This result shows the limits of differential analysis when this stops to the detection of DEGs without understanding how these are connected. For example *TNF*, target of drugs as infliximab for the treatment of CD [31], or some interleukins as *IL6* [18], would not have been individuated without a network vision of the DEGs found. The table 2 reports the genes associated to CD present in the module. The semantic similarities between the two lists, using the three roots of GO, was: 0.903 for CC, 0.842 for MF and 0.875 for BP. The mean of the three similarity measures was 0.873. This means that the lists are highly related in terms of GO terms.

**Table 2.** Genes associated with CD present in our module. They are reported the different names, the raw fold changes and classification in DEGs (1) and not DEGs (0).

| Entre | Gene Symbol | Gene Name | Raw | DE |
| --- | --- | --- | --- | --- |
| 842 | *CASP9* | caspase 9 | 1,02 | 0 |
| 920 | *CD4* | CD4 molecule | 2,06 | 0 |
| 930 | *CD19* | CD19 molecule | 2,99 | 1 |
| 1437 | *CSF2* | colony | 0,62 | 0 |
| 3122 | *HLA-DRA* | major | 15,8 | 1 |
| 3383 | *ICAM1* | intercellular | 5,95 | 1 |
| 3460 | *IFNGR2* | interferon gamma | 0,66 | 0 |
| 3553 | *IL1B* | interleukin 1, beta | 16,9 | 1 |
| 3569 | *IL6* | interleukin 6 | 6,59 | 0 |
| 3587 | *IL10RA* | interleukin 10 | 3,28 | 1 |
| 3606 | *IL18* | interleukin 18 | 1,52 | 0 |
| 3683 | *ITGAL* | integrin, alpha L | 2,88 | 1 |
| 3689 | *ITGB2* | integrin, beta 2 | 6,26 | 0 |
| 3717 | *JAK2* | Janus kinase 2 | 2,00 | 0 |
| 3845 | *KRAS* | v-Ki-ras2 Kirsten | 1,18 | 0 |
| 5777 | *PTPN6* | protein tyrosine | 1,37 | 0 |
| 5879 | *RAC1* | ras-related C3 | 1,58 | 0 |
| 6777 | *STAT5B* | signal transducer | 1,49 | 0 |
| 7124 | *TNF* | tumor necrosis | 1,96 | 0 |
| 7132 | *TNFRSF1A* | tumor necrosis | 1,15 | 0 |
| 7186 | *TRAF2* | TNF receptor- | 0,77 | 0 |
| 7535 | *ZAP70* | zeta-chain (TCR) | 4,55 | 0 |
| 8767 | *RIPK2* | Receptor serine- | 2,16 | 1 |
| 8807 | *IL18RAP* | interleukin 18 | 4,20 | 1 |
| 9564 | *LOC646079* | breast cancer | 0,75 | 0 |
| 10454 | *TAB1* | mitogen-activated | 0,49 | 0 |
| 22900 | *CARD8* | caspase | 1,92 | 1 |
| 114548 | *NLRP3* | NLR family, | 3,67 | 0 |

**Fig. 2.**
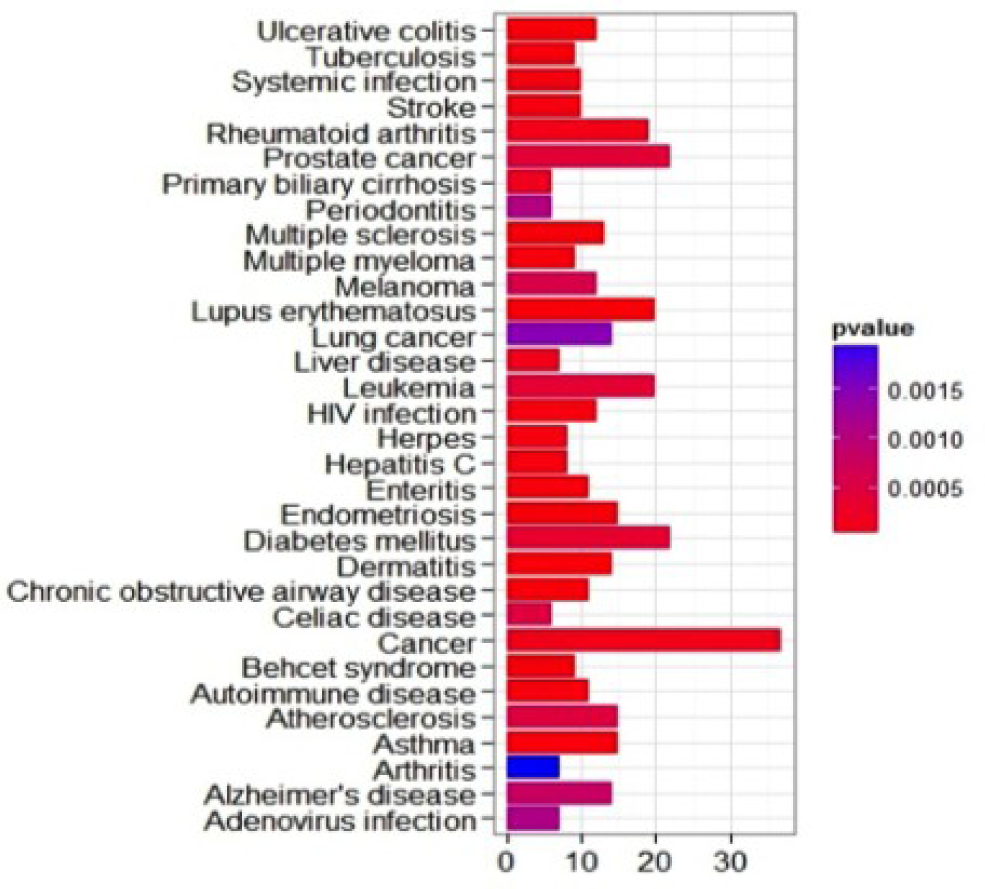
Diseases enriched by the genes in the module. To note the presence of diseases associated with CD as RA, UC and SLE.

#### 3.3 Topological analyses

The analysis of the topology of the network was finalized to find the most important genes in terms of centrality and connectivity. The following analyses were performed: 1) detection of the top 10 genes with the highest betweenness score and with the highest Bonacich’s power centrality measure (table 3); 2) detection of the cut vertices (table 4). The highest values of normalized betweenness were associated to the gene *RELA* or *P65* (0.24) and to the gene *PIK3R2*. The importance of the latter gene is noticed in literature. The p65 seems to play a key role in the regulation of intestinal inflammation of IBD having a profound proinflammatory activity. Furthermore, it was reported that the nuclear level of *P65* increased in lamina propria biopsy specimens from patients with CD in comparison with patients with UC and controls [32]. The gene *PIK3R2*, belonging to the family of the phosphatidylinositol 3-kinases (PI3-K), is involved in a broad range of cellular responses from cell cycle regulation, apoptosis, growth, and cell survival, making this a highly complex signaling network involved in cellular homeostasis. Dysregulation of this complex pathway can lead to diseases such as cancer, inflammation, and autoimmunity reactions, all associated with IBD [33]. Giving a look to the genes with highest Bonacich’s power index, the most relevant results are the TLR genes, *MY96* and the genes *CEBPB* and *CEBPG*. The TLR genes are Toll-like receptors (TLRs) and through the ligation on intestinal epithelial cells by bacterial products promotes epithelial cell proliferation, secretion of IgA into the gut lumen and expression of antimicrobial peptides. This establishes a microorganism-induced programme of epithelial cell homeostasis and repair in the intestine. Thus, dysregulated TLR signalling may impair commensal-mucosal homeostasis, critically contributing to acute and chronic intestinal inflammatory processes in IBD colitis and associated cancer [34]. *LY96* is a co-receptor of the *TLR4*, so it is involved in the same biological processes [35]. While the genes *CEBPB* and *CEBPG* are involved in the response to inflammation of intestinal epithelial cells [36].

**Table 3.** The genes with the highest betweenness and Bonacich’s power centrality measure. The entrez id, the gene symbol, the official gene name and the normalized betweennees value are reported.

| Entrez ID | Gene Symbol | Gene Name | Centrality | Type of centrality |
| --- | --- | --- | --- | --- |
| 836 | *CASP3* | caspase 3, apoptosis-related cysteine peptidase | 0,114 | Top betweenness |
| 2316 | *FLNA* | filamin A, alpha (actin binding protein 280) | 0,152 | Top betweenness |
| 3688 | *ITGB1* | integrin, beta 1 | 0,187 | Top betweenness |
| 5296 | *PIK3R2* | phosphoinositide-3-kinase, regulatory subunit 2 (beta) | 0,296 | Top betweenness |
| 5879 | *RAC1* | ras-related C3 botulinum toxin substrate 1 | 0,164 | Top betweenness |
| 5894 | *RAF1* | v-raf-1 murine leukemia viral oncogene homolog 1 | 0,163 | Top betweenness |
| 5970 | *RELA* | v-rel reticuloendotheliosis viral oncogene homolog A | 0,243 | Top betweenness |
| 6416 | *MAP2K4* | mitogen-activated protein kinase kinase 4 | 0,197 | Top betweenness |
| 6714 | *SRC* | v-src sarcoma, viral oncogene homolog | 0,122 | Top betweenness |
| 10451 | *VAV3* | vav 3 guanine nucleotide exchange factor | 0,134 | Top betweenness |
| 1051 | *CEBPB* | CCAAT/enhancer binding protein (C/EBP), beta | 2,356 | Top Bonacich power |
| 1054 | *CEBPG* | CCAAT/enhancer binding protein (C/EBP), gamma | 2,356 | Top Bonacich power |
| 4261 | *CIITA* | class II, major histocompatibility complex, transactivator | 2,313 | Top Bonacich power |
| 4802 | *NFYC* | nuclear transcription factor Y, gamma | 2,184 | Top Bonacich power |
| 5336 | *PLCG2* | phospholipase C, gamma 2 | 2,335 | Top Bonacich power |
| 5970 | *RELA* | v-rel reticuloendotheliosis viral oncogene homolog A | 2,277 | Top Bonacich power |
| 7097 | *TLR2* | toll-like receptor 2 | 2,320 | Top Bonacich power |
| 7099 | *TLR4* | toll-like receptor 4 | 2,320 | Top Bonacich power |
| 10673 | *TNFSF13B* | tumor necrosis factor (ligand) superfamily | 1,903 | Top Bonacich power |
| 23643 | *LY96* | lymphocyte antigen 96 | 2,363 | Top Bonacich power |

**Table 4.** Genes in the cut set of the CD modules.

| **Entrez ID** | **Gene Symbol** | **Gene Name** |
| --- | --- | --- |
| 5601 | *MAPK9* | mitogen-activated protein kinase 9 |
| 1857 | *DVL3* | dishevelled, dsh homolog 3 (Drosophila) |
| 6300 | *MAPK12* | mitogen-activated protein kinase 12 |
| 3606 | *IL18* | interleukin 18 (interferon-gamma-inducing factor) |
| 2932 | *GSK3B* | glycogen synthase kinase 3 beta |
| 8767 | *RIPK2* | receptor-interacting serine-threonine kinase 2 |
| 834 | *CASP1* | caspase 1, apoptosis-related cysteine peptidase (interleukin 1, beta, |
| 5530 | *PPP3CAA* | protein phosphatase 3 (formerly 2B), catalytic subunit, alpha isoform |
| 3695 | *ITGB7* | integrin, beta 7 |
| 7189 | *TRAF6* | TNF receptor-associated factor 6 |
| 4615 | *MYD88* | myeloid differentiation primary response gene (88) |
| 8174 | *MADCAM1* | mucosal vascular addressin cell adhesion molecule 1 |
| 2885 | *GRB2* | growth factor receptor-bound protein 2 |
| 7099 | *TLR4* | toll-like receptor 4 |
| 22900 | *CARD8* | caspase recruitment domain family, member 8 |
| 572 | *BAD* | BCL2-associated agonist of cell death |
| 5777 | *PTPN6* | protein tyrosine phosphatase, non-receptor type 6 |
| 6654 | *SOS1* | son of sevenless homolog 1 (Drosophila) |
| 114548 | *NLRP3* | NLR family, pyrin domain containing 3 |

The genes in the cut set, i.e, the genes which removal causes the separation of the module in more components, are the following: *MAPK9, MYD88, GRB2, IL18, CARD8, BAD, PTPN6, CASP1, NLRP3*. Most of these genes are also among the nodes with the highest value of centrality indexes. *IL18* was reported as a novel immunoregulatory cytokine with an important role in chronic inflammatory disorders as CD and it was proposed as new drug target so offers the theoretical advantage of affecting the inductive phase of Th1 cell activation [37]. Also the gene *PTPN6* or SHP-1 phosphates, a tumor suppressor gene and a master negative regulator of proinflammatory processes, may represent a new potential therapeutic targets for IBD [38]. Both the centrality measures and the articulation nodes had pointed out the relevance of central genes and articulation genes, in the comprehension of molecular mechanism of CD.

#### 3.4 Hierarchical analysis of significant shortest paths

As the 19 affected samples comes from 12 patients responding to infliximab treatment and from 7 not responding to infliximab, an hierarchical analysis on the significant shortest paths was done. Practically the idea was to look for, among the set of significant shortest paths, those that characterize the responsiveness to the infliximab treatment. On the 567 significant paths, 17 were associated to the responsiveness to the treatment, composed by 59 genes reported in Table 5.

**Table 5.** Genes in the shortest paths connected to the responsiveness to infliximab treatment in UC patients

| Entrez ID | Gene Symbol | Gene Name |
| --- | --- | --- |
| 1292 | *COL6A2* | collagen, type VI, alpha 2 |
| 1293 | *COL6A3* | collagen, type VI, alpha 3 |
| 6774 | *STAT3* | signal transducer and activator of transcription 3 |
| 5894 | *RAF1* | v-raf-1 murine leukemia viral oncogene homolog 1 |
| 2919 | *CXCL1* | chemokine (C-X-C motif) ligand 1 |
| 5578 | *PRKCA* | protein kinase C, alpha |
| 8767 | *RIPK2* | receptor-interacting serine-threonine kinase 2 |
| 7124 | *TNF* | tumor necrosis factor (TNF superfamily, member 2) |
| 59345 | *GNB4* | guanine nucleotide binding protein (G protein), beta polypeptide 4 |
| 5062 | *PAK2* | p21 protein (Cdc42/Rac)-activated kinase 2 |
| 4615 | *MYD88* | myeloid differentiation primary response gene (88) |
| 6416 | *MAP2K4* | mitogen-activated protein kinase kinase 4 |
| 84674 | *CARD6* | caspase recruitment domain family, member 6 |
| 8837 | *CFLAR* | CASP8 and FADD-like apoptosis regulator |
| 8324 | *FZD7* | frizzled homolog 7 (Drosophila) |
| 3579 | *CXCR2* | interleukin 8 receptor, beta |
| 3055 | *HCK* | hemopoietic cell kinase |
| 5601 | *MAPK9* | mitogen-activated protein kinase 9 |
| 1857 | *DVL3* | dishevelled, dsh homolog 3 (Drosophila) |
| 6885 | *MAP3K7* | mitogen-activated protein kinase kinase kinase 7 |
| 3716 | *JAK1* | Janus kinase 1 |
| 834 | *CASP1* | caspase 1, apoptosis-related cysteine peptidase |
| 836 | *CASP3* | caspase 3, apoptosis-related cysteine peptidase |
| 208 | *AKT2* | v-akt murine thymoma viral oncogene homolog 2 |
| 7132 | *TNFRSF1A* | tumor necrosis factor receptor superfamily, member 1A |
| 3717 | *JAK2* | Janus kinase 2 |
| 409 | *ARRB2* | arrestin, beta 2 |
| 1234 | *CCR5* | chemokine (C-C motif) receptor 5 |
| 3688 | *ITGB1* | integrin, beta 1 |
| 6777 | *STAT5B* | signal transducer and activator of transcription 5B |
| 3587 | *IL10RA* | interleukin 10 receptor, alpha |
| 83660 | *TLN2* | talin 2 |
| 5594 | *MAPK1* | mitogen-activated protein kinase 1 |
| 317 | *APAF1* | apoptotic peptidase activating factor 1 |
| 5970 | *RELA* | v-rel reticuloendotheliosis viral oncogene homolog A (avian) |
| 3606 | *IL18* | interleukin 18 (interferon-gamma-inducing factor) |
| 2932 | *GSK3B* | glycogen synthase kinase 3 beta |
| 7414 | *VCL* | vinculin |
| 3315 | *HSPB1* | heat shock 27kDa protein 1 |
| 7474 | *WNT5A* | wingless-type MMTV integration site family, member 5A |
| 2771 | *GNAI2* | guanine nucleotide binding protein (G protein), |
| 58484 | *NLRC4* | NLR family, CARD domain containing 4 |
| 9020 | *MAP3K14* | mitogen-activated protein kinase kinase kinase 14 |
| 3553 | *IL1B* | interleukin 1, beta |
| 3554 | *IL1R1* | interleukin 1 receptor, type I |
| 8717 | *TRADD* | TNFRSF1A-associated via death domain |
| 5829 | *PXN* | paxillin |
| 7855 | *FZD5* | frizzled homolog 5 (Drosophila) |
| 958 | *CD40* | CD40 molecule, TNF receptor superfamily member 5 |
| 1441 | *CSF3R* | colony stimulating factor 3 receptor (granulocyte) |
| 2920 | *CXCL2* | chemokine (C-X-C motif) ligand 2 |
| 122809 | *SOCS4* | suppressor of cytokine signaling 4 |
| 1398 | *CRK* | v-crk sarcoma virus CT10 oncogene homolog (avian) |
| 8807 | *IL18RAP* | interleukin 18 receptor accessory protein |
| 2829 | *XCR1* | chemokine (C motif) receptor 1 |
| 10454 | *TAB1* | mitogen-activated protein kinase kinase kinase 7 interacting protein |
| 7097 | *TLR2* | toll-like receptor 2 |
| 4772 | *NFATC1* | nuclear factor of activated T-cells, calcineurin-dependent 1 |
| 5296 | *PIK3R2* | phosphoinositide-3-kinase, regulatory subunit 2 (beta) |

### UC data

For the UC versus control samples, we found 2149 significant genes with a BH corrected p-value less than 0.01 and a minimum fold change of 1.5. Starting from DEGs we generated the relative module.

#### 3.1 Module generation

On the 2159 DEGs, SPIA revealed 22 significant KEGG pathways (see Table 6). Some of them were also found in CD as the cytokine-cytokine receptor interaction, the chemokine signaling pathway [17], the NF-▮B signaling pathway and the pathway of the CAMs. Other important pathways are the PI3K/Akt signal transduction pathway that is involved in the regulation and release of pro-inflammatory cytokines such as *TNF* and plays an important role in the development and progression of UC [39] and the NOD-like receptor pathway [40]. Most of the pathways had a positive accumulated perturbation except two of them, the PPAR signaling pathway and phosphatidylinositol signaling system, also if the pPERT was not significant. The pathways with the highest values of perturbation are the CAMs, the chemokines and the natural killer cell mediated cytotoxicity that have a clear function in the pathogenesis of UC [41].. The pathways were transformed in graph, merged. The shortest path procedure between each couple of DEGs gave as result 18752 shortest paths with 393 genes and 1411 edges involved. The significant shortest paths were 1586 out of 18752 involving 300 genes and 551 edges. Fig. 3 shows the graph obtained by the fusion of the 1586 significant paths.

**Table 6.** The KEGG perturbed pathways involved in UC. pSize represents the total number of nodes in the pathways, NDE the number of DEGs, tA the total perturbation, pPERT the p-value of the perturbation and pGFdr the adjusted p-value.

| Name | pSiz | ND | pND | tA | pPER | pGFd |
| --- | --- | --- | --- | --- | --- | --- |
| Cytokine-cytokine receptor interaction | 241 | 51 | 0,00 | 46,31 | 0,00 | 0,00 |
| Chemokine signaling pathway | 178 | 38 | 0,00 | 101,2 | 0,00 | 0,00 |
| Cell adhesion molecules (CAMs) | 88 | 25 | 0,00 | 63,23 | 0,02 | 0,00 |
| ECM-receptor interaction | 86 | 15 | 0,03 | 21,63 | 0,00 | 0,00 |
| B cell receptor signaling pathway | 73 | 20 | 0,00 | 13,70 | 0,32 | 0,00 |
| Natural killer cell mediated | 128 | 25 | 0,00 | 87,79 | 0,01 | 0,00 |
| PPAR signaling pathway | 63 | 17 | 0,00 | -1,68 | 0,35 | 0,01 |
| Small cell lung cancer | 82 | 19 | 0,00 | 25,51 | 0,10 | 0,01 |
| Apoptosis | 84 | 21 | 0,00 | 6,52 | 0,69 | 0,01 |
| NOD-like receptor signaling pathway | 48 | 13 | 0,00 | 14,76 | 0,14 | 0,01 |
| Amoebiasis | 46 | 13 | 0,00 | 8,33 | 0,24 | 0,01 |
| NF-kappa B signaling pathway | 75 | 17 | 0,00 | 20,44 | 0,12 | 0,02 |
| Legionellosis | 39 | 11 | 0,00 | 9,70 | 0,21 | 0,03 |
| Tuberculosis | 161 | 307 | 0,00 | 13,04 | 0,48 | 0,03 |
| Toxoplasmosis | 91 | 19 | 0,00 | 11,38 | 0,30 | 0,04 |
| Phosphatidylinositol signaling system | 76 | 15 | 0,01 | -2,25 | 0,08 | 0,04 |
| HTLV-I infection | 198 | 33 | 0,00 | 20,09 | 0,23 | 0,04 |
| Staphylococcus aureus infection | 26 | 8 | 0,00 | 14,01 | 0,23 | 0,04 |
| Focal adhesion | 203 | 30 | 0,03 | 44,82 | 0,03 | 0,04 |
| T cell receptor signaling pathway | 97 | 20 | 0,00 | 7,71 | 0,55 | 0,05 |
| PI3K-Akt signaling pathway | 340 | 49 | 0,01 | 41,56 | 0,12 | 0,05 |
| Malaria | 11 | 5 | 0,00 | 2,64 | 0,35 | 0,05 |

**Fig. 3.**
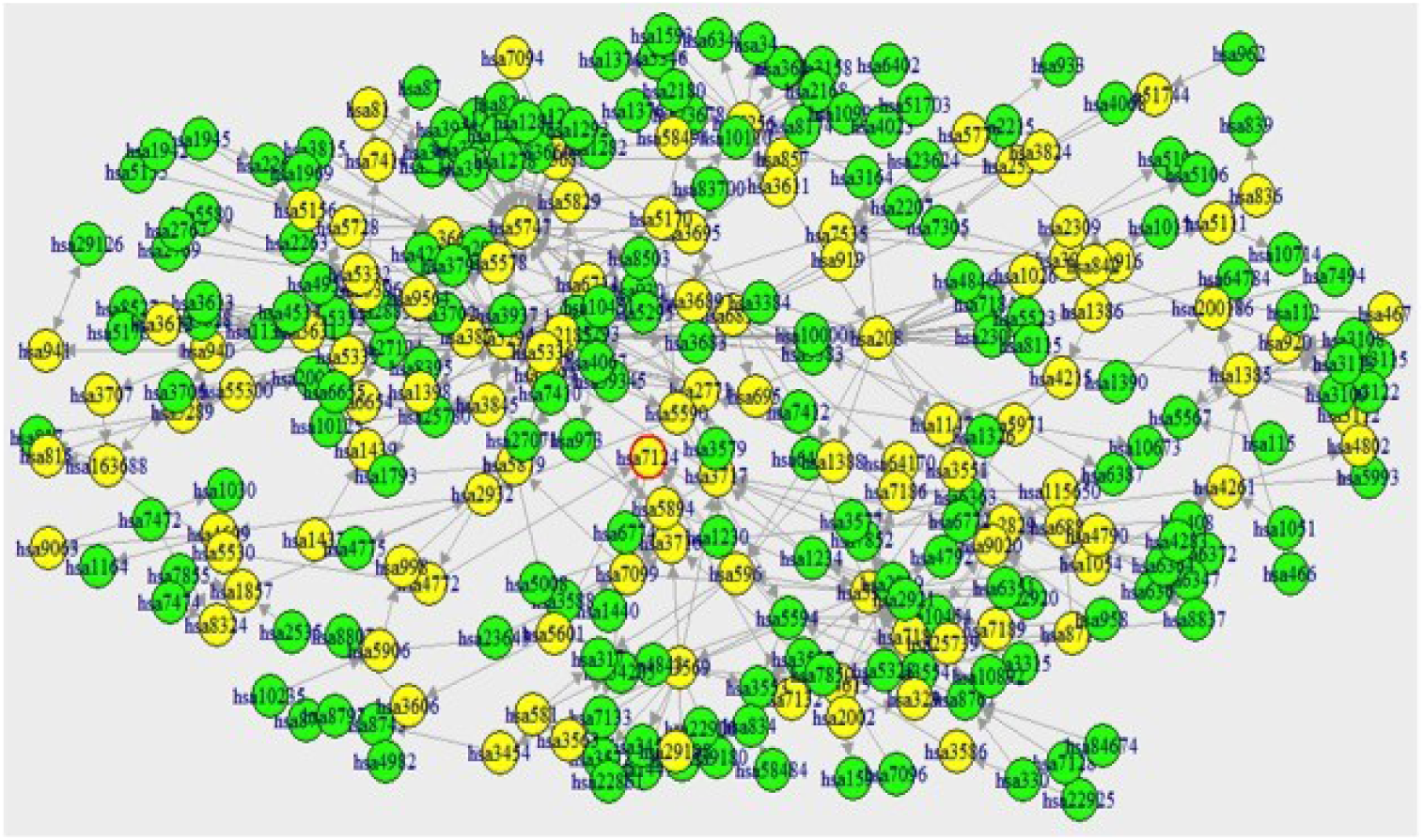
Module obtained by merging the significant shortest paths between DEGs found for UC samples. Green nodes are DEGs while yellow are not DEGs. Also here the gene 7124 circled in red was not found as DEG and it would not have been considered without the network approach.

**Fig. 4.**
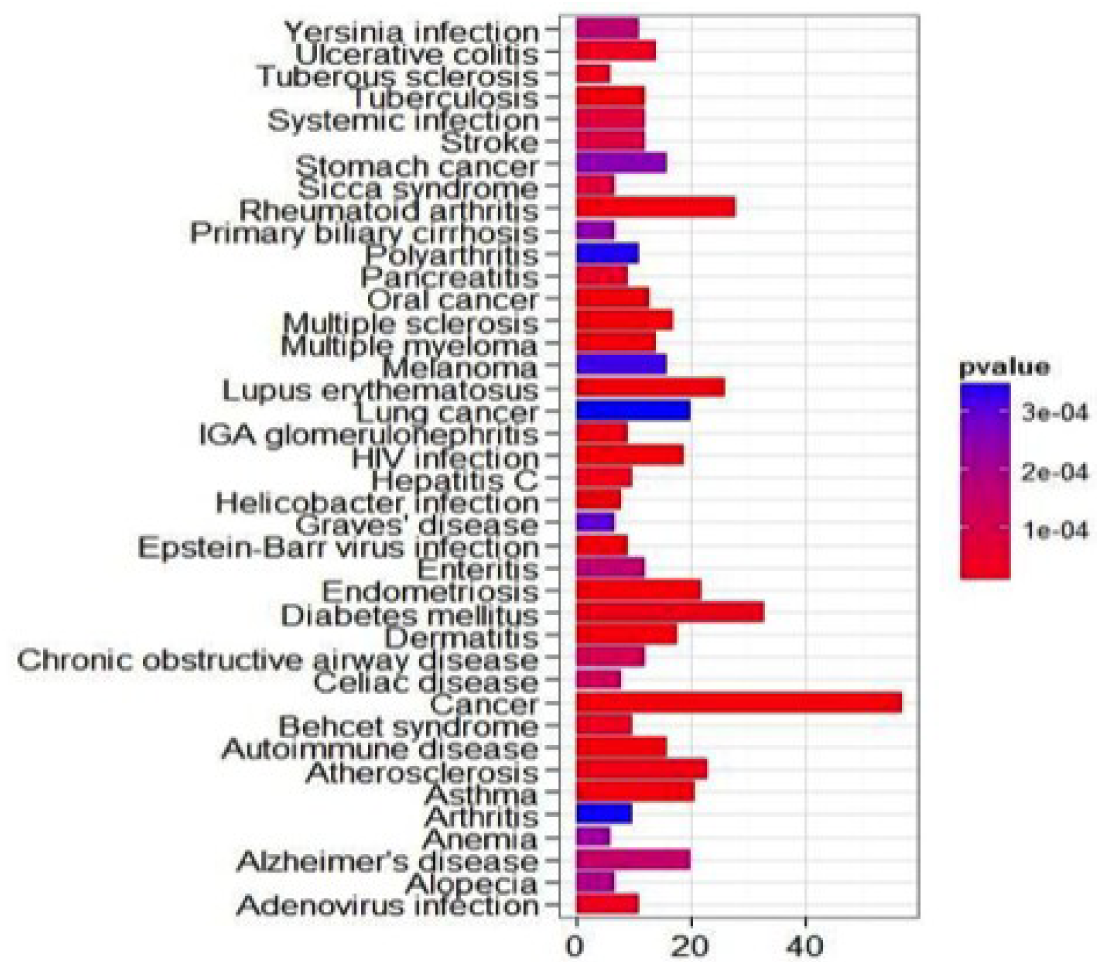
Diseases enriched by the UC genes in the module. To note the presence of UC. This means that the selected genes in the module are statistically associated with UC

#### 3.2 Module validation

For the validation of the module we performed two different analyses as the previous example. The result of the DO enrichment analysis confirms the goodness of the procedure considering that UC was found among the enriched diseases and other diseases that involves the same molecular mechanisms as Yersinia infection, SLE, RA, hepatitis were found. For semantic similarity validation, we looked for the genes associated with UC in the database “Gene” of NCBI. We individuated 461 unique Entrez identifiers. The intersection of the genes in the module with those associated to UC revealed 44 genes in common. Of these 25 were DEGs while the remaining genes were not DEGs. Although *TNF* and *RELA* are important genes for pathogenesis of UC, they were not among DEGs, but our procedure allowed to characterize them. Table 7 reports the genes associated to UC present in the module. The semantic similarities between the two lists, using the three roots of GO, was: 0.912 for CC, 0.844 for MF and 0.894 for BP. The mean of the three similarity measures was 0.883.

**Table 7.** Genes associated with UC present in our module. The different names, the raw fold changes and classification in DEGs (1) and not DEGs (0) are reported.

| Entrez | Gene symbol | Gene Name | Raw FC | DEG (1/0) |
| --- | --- | --- | --- | --- |
| 330 | BIRC3 | baculoviral IAP repeat-containing 3 | 5,45 | 1 |
| 842 | CASP9 | caspase 9 | 1,04 | 0 |
| 941 | CD80 | CD80 molecule | 2,56 | 0 |
| 958 | CD40 | CD40 molecule | 16,68 | 1 |
| 2215 | FCGR3B | Fc fragment of IgG | 6,17 | 1 |
| 3122 | HLA-DRA | major histocompatibility complex, class II, DR alpha | 7,84 | 1 |
| 3383 | ICAM1 | intercellular adhesion molecule 1 | 4,89 | 1 |
| 3553 | IL1B | interleukin 1, beta | 4,17 | 1 |
| 3569 | IL6 | interleukin 6 (interferon, beta 2) | 3,65 | 0 |
| 3575 | IL7R | interleukin 7 receptor | 0,60 | 1 |
| 3577 | CXCR1 | interleukin 8 receptor, alpha | 1,43 | 1 |
| 3579 | CXCR2 | interleukin 8 receptor, beta | 3,45 | 1 |
| 3586 | IL10 | interleukin 10 | 1,91 | 0 |
| 3587 | IL10RA | interleukin 10 receptor, alpha | 1,10 | 1 |
| 3588 | IL10RB | interleukin 10 receptor, beta | 8,64 | 1 |
| 3606 | IL18 | interleukin 18 (interferon-gamma-inducing factor) | 4,79 | 0 |
| 3683 | ITGAL | integrin, alpha L | 0,68 | 1 |
| 3717 | JAK2 | Janus kinase 2 | 1,06 | 0 |
| 3845 | KRAS | v-Ki-ras2 Kirsten rat sarcoma viral oncogene homolog | 0,86 | 0 |
| 4023 | LPL | lipoprotein lipase | 1,44 | 1 |
| 4261 | CIITA | class II, major histocompatibility complex, transactivator | 1,41 | 0 |
| 4772 | NFATC1 | nuclear factor of activated T-cells | 2,37 | 0 |
| 4790 | NFKB1 | nuclear factor of kappa light polypeptide gene enhancer in B-cells 1 | 1,93 | 0 |
| 4843 | NOS2 | nitric oxide synthase 2, inducible | 6,30 | 1 |
| 5728 | PTENP1 | phosphatase and tensin homolog | 1,89 | 0 |
| 5777 | PTPN6 | protein tyrosine phosphatase, non-receptor type 6 | 1,25 | 0 |
| 5970 | RELA | v-rel reticuloendotheliosis viral oncogene homolog A | 2,17 | 0 |
| 6347 | CCL2 | chemokine (C-C motif) ligand 2 | 1,02 | 1 |
| 6772 | STAT1 | signal transducer and activator of transcription 1, 91kDa | 2,91 | 1 |
| 6774 | STAT3 | signal transducer and activator of transcription 3 | 4,95 | 1 |
| 7099 | TLR4 | toll-like receptor 4 | 0,42 | 0 |
| 7124 | TNF | tumor necrosis factor (TNF superfamily, member 2) | 9,80 | 0 |
| 7132 | TNFRSF1A | tumor necrosis factor receptor superfamily, member 1A | 3,05 | 0 |
| 7133 | TNFRSF1B | tumor necrosis factor receptor superfamily, member 1B | 0,64 | 1 |
| 7186 | TRAF2 | TNF receptor-associated factor 2 | 1,73 | 0 |
| 7494 | XBP1 | X-box binding protein 1 | 8,83 | 1 |
| 7535 | ZAP70 | zeta-chain (TCR) associated protein kinase 70kDa | 2,07 | 0 |
| 7850 | IL1R2 | interleukin 1 receptor, type II | 5,37 | 1 |
| 7852 | CXCR4 | chemokine (C-X-C motif) receptor 4 | 0,37 | 1 |
| 8174 | MADCAM1 | mucosal vascular addressin cell adhesion molecule 1 | 0,70 | 1 |
| 8743 | TNFSF10 | tumor necrosis factor (ligand) superfamily, member 10 | 6,37 | 1 |
| 8837 | CFLAR | CASP8 and FADD-like apoptosis regulator | 0,89 | 1 |
| 22900 | CARD8 | caspase recruitment domain family, member 8 | 4,58 | 1 |
| 64170 | CARD9 | caspase recruitment domain family, member 9 | 1,33 | 0 |

#### 3.3 Topological analyses

The analysis of the topology of the network was finalized to find the most important genes in terms of centrality and connectivity. The following analyses were performed: 1) detection of the top 10 genes with the highest betweenness score and with highest Bonacich’s power centrality measure (table 8); 2) detection of the cut vertices (table 9). The highest betweenness value ara associated to the gene *PIK3R2* (0.26) and to the gene *AKT2* (0.28). The gene *PIK3R2* was also found for CD and its involvedment in IBD was previously described [33]. About the gene *AKT2*, it was shown that the expression level of the AKT pathway was increased in UC patients and the treatment with wortmanninacting on this gene improved the health of the patients [39]. About the Bonacich’s power the highest value was for *IL6* (3.84) a proinflammatory cytokine that has been detected in both colonic biopsy specimens and serum samples of patients with UC which levels are correlated with the disease severity [42]. When we look to the cut set geven, genes as *TNF*, interleukines, *TLR* were included. All of them are key genes in the pathogenesis of UC.

**Table 8.** The genes with the highest betweenness and Bonacich’s power centrality measure. The entrez id, the gene symbol, the official gene name and the normalized betweennees value are reported.

| Genes | Gene Symbol | Gene Name | Centrality Measure | Type of centrality |
| --- | --- | --- | --- | --- |
| 208 | AKT2 | v-akt murine thymoma viral oncogene homolog 2 | 0,26 | Top betweenness |
| 1147 | CHUK | conserved helix-loop-helix ubiquitous kinase | 0,13 | Top betweenness |
| 2829 | XCR1 | chemokine (C motif) receptor 1 | 0,12 | Top betweenness |
| 3688 | ITGB1 | integrin, beta 1 | 0,10 | Top betweenness |
| 3717 | JAK2 | Janus kinase 2 | 0,14 | Top betweenness |
| 5296 | PIK3R2 | phosphoinositide-3-kinase, regulatory subunit 2 | 0,28 | Top betweenness |
| 5879 | RAC1 | ras-related C3 botulinum toxin substrate 1 | 0,09 | Top betweenness |
| 5970 | RELA | v-rel reticuloendotheliosis viral oncogene | 0,10 | Top betweenness |
| 5971 | RELB | v-rel reticuloendotheliosis viral oncogene | 0,13 | Top betweenness |
| 6363 | CCL19 | chemokine (C-C motif) ligand 19 | 0,11 | Top betweenness |
| 1440 | CSF3 | colony stimulating factor 3 (granulocyte) | 2,07 | Top Boncich power |
| 3569 | IL6 | interleukin 6 (interferon, beta 2) | 3,84 | Top Boncich power |
| 3586 | IL10 | interleukin 10 | 2,21 | Top Boncich power |
| 3587 | IL10RA | interleukin 10 receptor, alpha | 2,07 | Top Boncich power |
| 3588 | IL10RB | interleukin 10 receptor, beta | 2,07 | Top Boncich power |
| 3716 | JAK1 | Janus kinase 1 | 1,92 | Top Boncich power |
| 5008 | OSM | oncostatin M | 2,07 | Top Boncich power |
| 6772 | STAT1 | signal transducer and activator of transcription 1, | 1,87 | Top Boncich power |
| 22925 | PLA2R1 | phospholipase A2 receptor 1, 180kDa | 2,36 | Top Boncich power |
| 1051 | CEBPB | CCAAT/enhancer binding protein (C/EBP), beta | 1,87 | Top Boncich power |

**Table 9.**
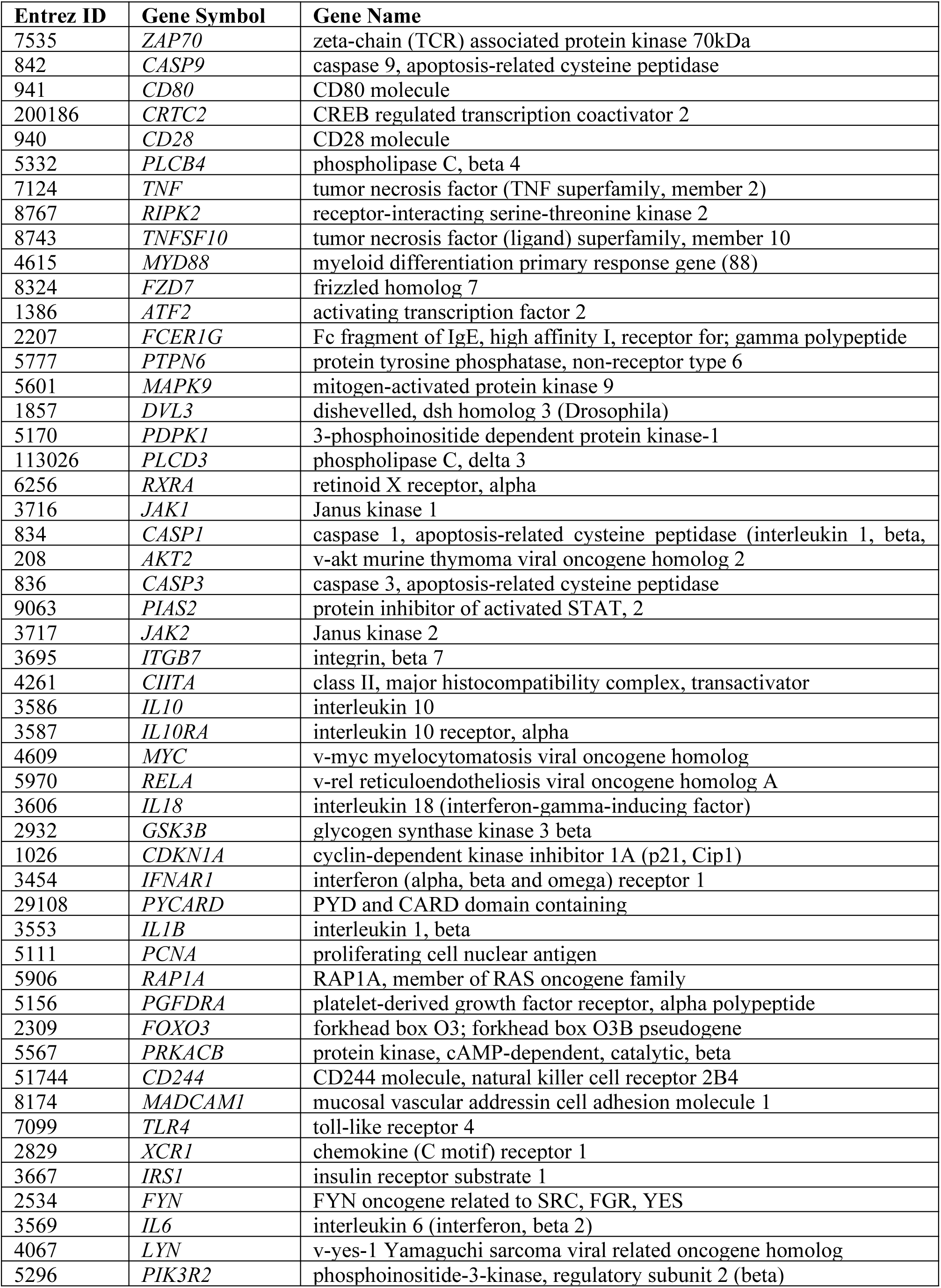
Genes in the cut set of the UC modules.

#### 3.4 Hierarchical analysis of significant shortest paths

As the 24 UC samples comes from 8 patients responding to infliximab treatment and 16 not responding to infliximab, an hierarchical analysis was performed on the significant shortest paths between responding against not responding.

On the 1586 significant paths, only 3 were associated to the responsiveness to the treatment, composed by 19 genes reported in the Table 10.

**Table 10.**
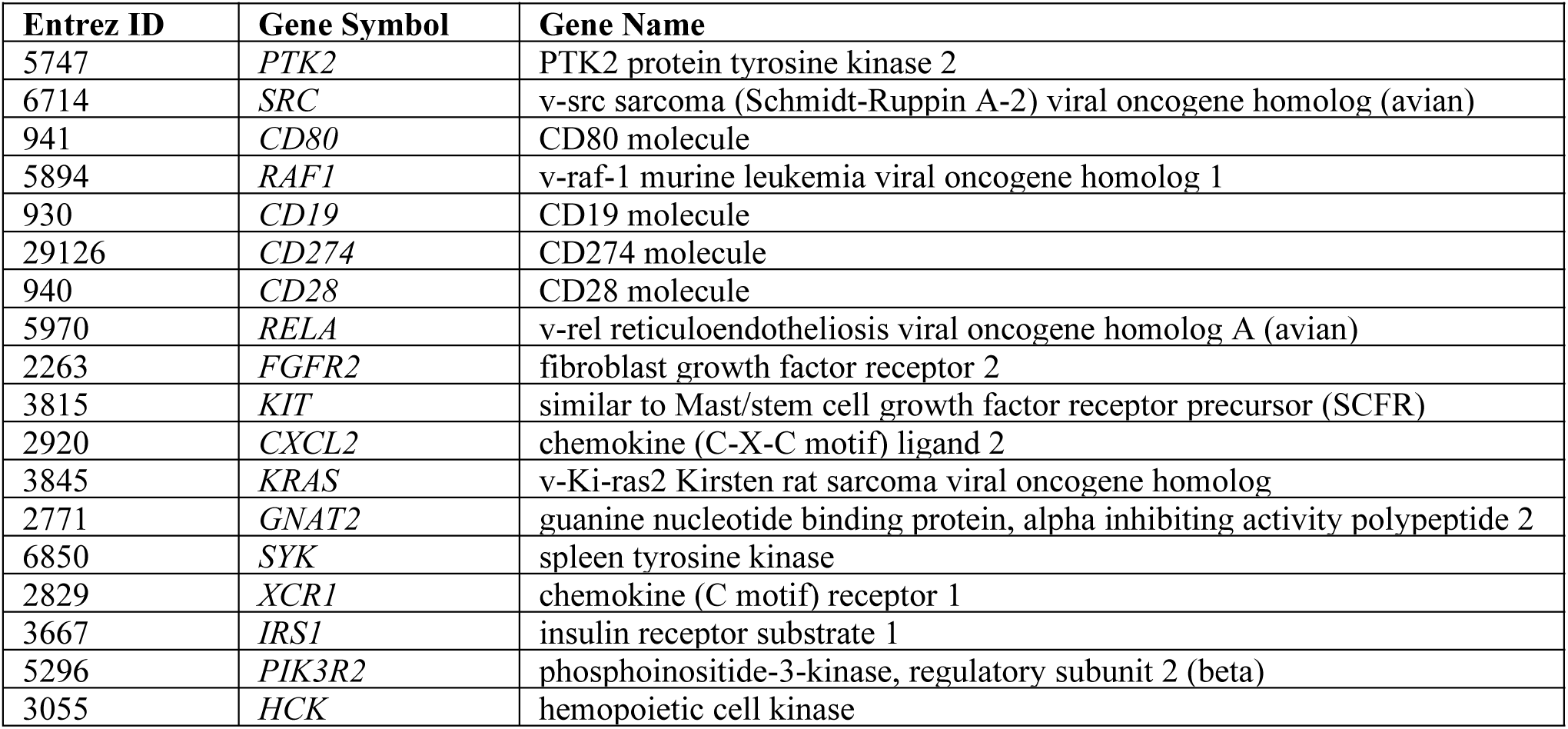
Genes found in the significant shortest paths for the responsiveness to infliximab treatment in UC patients.

| Entrez ID | Gene Symbol | Gene Name |
| --- | --- | --- |
| 5747 | *PTK2* | PTK2 protein tyrosine kinase 2 |
| 6714 | *SRC* | v-src sarcoma (Schmidt-Ruppin A-2) viral oncogene homolog (avian) |
| 941 | *CD80* | CD80 molecule |
| 5894 | *RAF1* | v-raf-1 murine leukemia viral oncogene homolog 1 |
| 930 | *CD19* | CD19 molecule |
| 29126 | *CD274* | CD274 molecule |
| 940 | *CD28* | CD28 molecule |
| 5970 | *RELA* | v-rel reticuloendotheliosis viral oncogene homolog A (avian) |
| 2263 | *FGFR2* | fibroblast growth factor receptor 2 |
| 3815 | *KIT* | similar to Mast/stem cell growth factor receptor precursor (SCFR) |
| 2920 | *CXCL2* | chemokine (C-X-C motif) ligand 2 |
| 3845 | *KRAS* | v-Ki-ras2 Kirsten rat sarcoma viral oncogene homolog |
| 2771 | *GNAT2* | guanine nucleotide binding protein, alpha inhibiting activity polypeptide 2 |
| 6850 | *SYK* | spleen tyrosine kinase |
| 2829 | *XCR1* | chemokine (C motif) receptor 1 |
| 3667 | *IRS1* | insulin receptor substrate 1 |
| 5296 | *PIK3R2* | phosphoinositide-3-kinase, regulatory subunit 2 (beta) |
| 3055 | *HCK* | hemopoietic cell kinase |

## Conclusions

The reductionist method of dissecting biological systems in smaller components has had its importance in the explanation of numerous biological processes. The most of biological data analyses were influenced from this point of view. The classical gene expression data analysis supplied a list of DEGs but it was not able to explain the complex molecular mechanisms behind the observed DEGs. It is clear that the specificity of a complex biological activity does not arise from the specificity of the individual genes that are involved, but results from the way in which the genes interact for absolving specific biological functions. Recent progress in the field of module level analysis promises to revolutionize our view of systems biology and the relative analysis approaches. In this paper we proposed a module level analysis for gene expression data that could take over the present methods for the individuation of modules. In fact it is used to divide the individuation approaches in network-based, expression-based and pathway based. Our approach is a mixed approach that starts from perturbed pathways in which the DEGs are involved, for individuating the significant shortest paths using network and expression information. The new concept is surely connected to the use of SEM for testing the significance of each shortest path model between the groups considered in the experiment. The procedure was tested on a microarray finalized to unravel the molecular mechanisms of diseases as CD and UC. We took the data relative to the gene expression from samples of CD and UC patients. After the t-test and pathway analysis with SPIA, the significant pathways were merged in an unique graph. Then all the shortest paths, that connect DEGs by other DEGs or other microarray genes, were found. The differential analysis of the shortest paths revealed significant shortest paths that characterize the experimental group on the control. The module obtained merging all the significant shortest paths allowed to detect the key molecular network that could explain the disease. The importance of this approach is demonstrated from the fact that very important genes associated to CD and UC, as the *TNF* or many interleukins, would not been individuated without the comprehension of the interactions between the DEGs. The validation of the module by enrichment analysis on DO has highlighted that the genes present in the module have an important role in the progression and characterization of many diseases with a common molecular root with CD and UC, as RA and SLE. Furthermore using the Wang similarity semantic index based on GO structure, we found that the module genes and the genes associate “a priori” with CD and UC have high values. Finally the network analysis on the modules, finalized to discover the most central genes and the genes fundamental for the connectivity of the module allowed to reveal possible markers for the diseases. The betweenness and the Bonacich’s power centrality measures revealed that the most central genes (*P65, MYD88, IL6*) are key genes in the inflammatory processes associated principally with chemokine, cytokine and NF-kappa B signaling pathways. For the connectivity, most of the articulation nodes are also the most central nodes, but it is noteworthy the gene *IL18*, proposed as possible drug target for diseases as the CD. In conclusion our approach, in basis to the results obtained, is surely notable as new approach of the downstream analysis of gene expression data. Surely, it could be applied also to RNA-seq data and improved considering also the neighbors that directly influences the DEGs.

### Competiting interests

The author declares having no competing interests.

